# Failure of *in vitro* differentiation of *Plasmodium falciparum* gametocytes into ookinetes arises because of poor gamete fertilisation

**DOI:** 10.1101/216721

**Authors:** Michael J Delves, Sara R Marques, Andrea Ruecker, Ursula Straschil, Celia Miguel-Blanco, María J López-Barragá, Joël Lelièvre, Irene Molina, Melanie Wree, Shinji L Okitsu, Elizabeth Winzeler, Fengwu Li, Joseph Vinetz, Sam Sheppard, Joana Guedes, Nadia Guerra, Esperanza Herreros, Robert E Sinden, Jake Baum

**Author notes:** These authors contributed equally to this work (Michael J Delves, Sara R Marques). Correspondence to: Dr Michael Delves and Dr Jake Baum, Department of Life Sciences, Imperial College London, Exhibition Road, South Kensington, London SW7 2AZ, United Kingdom.

## Abstract

A critical step towards malaria elimination will be the interruption of *Plasmodium* transmission from the human host to the mosquito. At the core of the transmission cycle lies *Plasmodium* sexual reproduction leading to zygote formation and mosquito midgut colonisation by ookinetes. Whilst *in vitro* ookinete culture from the murine and avian malaria parasites, *Plasmodium berghei* and *P. gallinaceum*, has greatly increased our knowledge of transmission biology; efforts to mimic the process in the human parasite *P. falciparum* have, to date, had only limited success. Using fluorescence microscopy and flow cytometry with antibodies specific to the male gametocyte and developing ookinetes, we sought to evaluate *P. falciparum* ookinete production using previously published *in vitro* protocols. We then compared *in vitro* versus *in vivo* ookinete production in both *P. falciparum* and *P. berghei* parasites, exploring potential barriers to complete development. Finally, we sought to test a wide range of literature-led culture conditions towards further optimisation of *in vitro P. falciparum* ookinete production. Despite extensive testing, our efforts to replicate published methods did not produce appreciable quantities of fully formed *P. falciparum* ookinetes *in vitro*. In parallel, however, gametocyte cultures that failed to differentiate fully *in vitro* successfully developed into ookinetes *in vivo* with an efficiency approximating that of *P. berghei*. Flow cytometry analysis showed that this disparity likely lies with the poor fertilization of *P. falciparum* gametes *in vitro*. Attempts to improve gametocyte fertility or define conditions more permissive to fertilisation/ookinete survival *in vitro* were also unsuccessful. Current *in vitro* conditions for *P. falciparum* ookinete production are not optimal for gamete fertilisation either due to the lack of parasite-species-specific mosquito factors missing from *in vitro* culture, or non-permissive cues contaminating culture preparations.

## 1. Introduction

The development and evaluation of reliable transmission blocking vaccines or drugs against human malaria, relies on our ability to study the sexual biology of the parasites. Sexual development underpins this passage of the parasites from man to mosquito with transmission itself mediated by mature gametocytes, formed in the vertebrate host but only reaching full maturity with production of gametes when ingested during a mosquito feed. Within the first hour in the vector midgut, gametes fertilize forming zygotes and over the ensuing 24 to 36 hours develop into motile and invasive ookinetes. Ookinete development is a major bottleneck of the parasite life-cycle and a priority transmission blocking-target (Sinden, 2010). Most of our current understanding of ookinete biology comes from data generated in *Plasmodium berghei* and *P. gallinaceum*, rodent and avian malaria parasites, respectively (Angrisano et al., 2012; Gass, 1979; Mehlhorn et al., 1980). First reports of *P. berghei* ookinete development *in vitro* date back to the 1960’s (Alger, 1968; Rosales-Ronquillo and Silverman, 1974; Yoeli and Upmanis, 1968) and were initially difficult to reproduce (Weiss and Vanderberg, 1977). These production protocols were significantly improved in the following decades (C J Janse et al., 1985; C. J. Janse et al., 1985; Sinden et al., 1985), and now permit both high-throughput (Delves et al., 2012) and high volume (Lal et al., 2009) culture options. In contrast, reports of ookinete development in the human malaria parasite *P. falciparum* appeared much later and described only limited success, revealing possible differences in the metabolic organisation of the mosquito stages between parasite species (Bounkeua et al., 2010; Carter et al., 1987; Ghosh et al., 2010).

Our goal here was to understand where the inefficiencies lie in previously published protocols for *in vitro P. falciparum* ookinete production. We tested a range of different conditions for *P. falciparum* cultures and compared sexual development in both *P. berghei* and *P. falciparum* by assessing their fertilization rates and capacity to develop into infectious ookinetes, in culture and in the mosquito midgut.

## 2. Materials and Methods

### 2.1. Ethics Statement

All procedures were performed in accordance with the UK Animals (Scientific Procedures) Act (PPL 70/7185) and approved by the Imperial College Animal Welfare and Ethical Review Body (AWERB).

### 2.2. Parasite production, culture and mosquito infection

The following parasite lines were used: *P. berghei* strain ANKA 2.34, PbGESTKO (Talman et al., 2011) and *P. falciparum* NF54 and 3D7 strains. *P. berghei* strains were a kind gift from R.E. Sinden (Imperial College London) and *P. falciparum* strains were a kind gift from R. Dinglasan (University of Florida) and the late David Walliker (University of Edinburgh). Equivalent strains are freely available through BEI Resources, NIAID, NIH (www.beiresources.org) or European Malaria Reagent Repository (www.malariaresearch.eu). *P. berghei* was maintained by cyclic passage in 6-to 8-week-old female Tucks Ordinary (TO) mice. *P. berghei* gametocyte and ookinete production, conversion rate determination and *Anopheles stephensi* infection were performed as previously described (Marques et al., 2014). *P. falciparum* asexual parasite maintenance and gametocyte growth were performed as previously described (Delves et al., 2016) using modifications defined. Unless otherwise stated, NF54 strain was used for all experiments. The ability of gametocytes to form gametes was confirmed by monitoring male gametogenesis (exflagellation) before attempting ookinete culture or mosquito feeds. A gametocyte culture was deemed fertile if more than 50 exflagellation centres per field of view were observed (using a 10x magnification objective) 20 minutes after activation, in a preparation where erythrocytes formed a tight monolayer. Cultured *P. falciparum* gametocytes were activated to form gametes by a decrease in temperature and addition of xanthurenic acid-containing ookinete medium (prepared as previously described (Bounkeua et al., 2010)) at 14 to 18 days post-induction.

*P. falciparum* ookinete cultures were performed as previously described (Bounkeua et al., 2010; Ghosh et al., 2010). Both protocols yielded comparably low conversion rates. Equivalent rates of conversion were seen when simplifying previous protocols to a generic, simplified protocol with the following modifications: Briefly, gametocyte cultures were spun down at 37°C and 300 rcf in 15 ml tubes and spent medium removed. A volume of human serum (Interstate Bloodbank) equal to the infected erythrocyte pellet was added at room temperature (approximately 24°C), diluting the hematocrit to 50%. Cells were then resuspended by gently flicking the tubes. 30 minutes after activation 5 volumes of ookinete medium (RPMI-1640, 25mM HEPES, 2mM L-glutamine, 2g/L sodium bicarbonate, 50mg/L hypoxanthine, pH 8.2) (Bounkeua et al., 2010) at room temperature were added to the culture, mixed by flicking, and then incubated at 26°C for 26hrs. Conversion rates were estimated as explained below. Numerous modifications to the previously published conditions were systematically tested - see Supplementary Tables 1+2 for details. Results of these modifications are described in Fig. 5 and Supplementary Tables 1+2.

For *Anopheles coluzzii* N’gousso strain *(Anopheles gambiae* M form) infections with *P. falciparum*, gametocyte cultures were spun down as above and spent medium was removed. Initial gametocytaemia in the cultures was reduced to 0.3% with uninfected blood at 50% hematocrit. Blood/parasite cell pellets were resuspended with equal volumes of pre-warmed human serum and transferred to membrane feeders. 40-50 mosquitoes were allowed to feed for 30 min. Mosquitoes were maintained on fructose at a constant temperature of 26°C and relative humidity of 80% until midgut dissection 26hrs later.

### 2.3. Live parasite detection and imaging

Twenty-six hours post-activation (hpa), cultured parasites were spun down, re-suspended in the appropriate staining solution and transferred to slides. For midgut-derived parasites, the blood bolus was removed from the midgut, transferred to glass slides, dispersed in ookinete medium and stained. Parasites were incubated for 20 minutes with Cy3-conjugated mouse monoclonal antibodies 4B7 for *P. falciparum* (obtained through BEI Resources NIAID, NIH: Monoclonal Antibody 4B7 Anti-*Plasmodium falciparum* 25 kDa Gamete Surface Protein (Pfs 25), MRA-28, contributed by David C. Kaslow) or 13.1 (Tirawanchai and Sinden, 1990) for *P. berghei* (kindly provided by R. E. Sinden, Imperial College London), both at a concentration of 1:300 in ookinete medium containing Hoechst 33342 at a concentration of 1 μg/μl; thus identifying female gamete-ookinete specific proteins Pfs25 and Pbs28, respectively. Parasite samples were visualized without washing on a Leica DMR microscope, and imaged with the Zeiss AxioCam HRC and Axiovision software (release 4.8 software). For Giemsa staining of midgut parasites, the blood bolus was removed and dispersed on a glass as above before being air-dried on a slide warmer. Slides were then fixed briefly in methanol and stained for 15 min in Giemsa stain. Ookinetes were then imaged by light microscopy as above.

### 2.4. Ookinete stage classification and conversion rate estimation

Both *in vivo* and *in vitro*, the morphological progression of sexual development for *P. berghei* ookinetes is similar consistent with observations on sexual development published previously (C J Janse et al., 1985). Gametes are formed within 20 minutes of activation, fertilization occurs within the first hour, and zygote to ookinete development ensues in the next 24 - 36 hours. Zygotes or stage I ookinetes are round or oval shaped. Stage II ookinetes, display small apical protrusions without pigment (3-6 hours) and stage III ookinetes (6-8 hours) have pigmented protrusions, both of these are termed retorts. Stage IV ookinetes have a round residual body and at stage V the round nucleus has migrated from the posterior to the middle of the retort extension. Conversion rates were calculated as the percentage of female gametes that convert into retorts/ookinetes. At least 500 cultured females/retorts were manually counted at a 40X objective magnification for each experimental replicate. For mosquito infections, at least 100 females/retorts/ookinetes were counted in each midgut of at least 12 mosquitoes for each experimental replicate.

### 2.5. Flow cytometry

Parasites were incubated for 20 minutes with antibody and Hoechst 33342, as above, washed twice in PBS, and fixed for 20 minutes in 4% paraformaldehyde (PFA). Cy3 and Hoechst fluorescence was analysed by flow cytometry using a BD LSR Fortessa cytometer. At least 10,000 cells were analysed per sample in three biological replicates. Data analysis was performed using FlowJo software v10.

### 2.6. Antibody generation

Anti-2606 was generated against the *P. falciparum* protein PF3D7_1325200, identified by analysis of its transcription profile, showing this *P. falciparum* putative oxidoreductase as being highly expressed in gametocytes (http://plasmodb.org/plasmo/). To produce polyclonal serum, peptides (comprising MTSVKHPKISVLGAGDIGC-Amide) were immunized into rabbits with serum affinity purified using standard procedures. Antibody generation and purification was performed commercially by 21st Century Biochemicals.

Anti-2606 antibody recognizes a protein of the predicted size of the oxidoreductase by Western blot and strongly detects both a subpopulation of non-activated gametocytes, likely male based on sex ratio (Supplementary Fig. 1) and exflagellating male gametocytes and gametes.

### 2.7. Immunocytochemistry and imaging

Smeared parasite samples were fixed at different time points in 4% PFA for 20 minutes. Rabbit polyclonal anti-2606 1:500 was used in combination with 4B7-Cy3. Alexa 488-conjugated anti-rabbit IgG for fluorescence detection was used at 1:1000 (A32731, ThermoFisher Scientific). Slides were mounted in Vectashield with DAPI (Vector Labs). Parasites were visualized using a Nikon Eclipse Ti inverted microscope, and images were analysed with NIS-Elements software v3.0.

### 2.8. Statistical analysis

Statistical analysis was performed with Graphpad Prism v7 using the unpaired student’s T-test.

## 3. Results

### 3.1. P. falciparum ookinete development in vitro is abnormal and/or fails to complete

*P. berghei* ookinete conversion rates in live cells is routinely measured using 13.1-Cy3, a fluorophore-conjugated antibody that recognizes ookinete surface antigen Pbs28 (Reininger et al., 2009). To achieve equivalent *P. falciparum* live cell detection, we used 4B7-Cy3, a monoclonal antibody that recognizes Pfs25 (Ruecker et al., 2014). This Pbs28 ortholog is similarly found at the surface of *P. falciparum* females, zygotes, retorts and ookinetes (Vermeulen et al., 1985) and allows live detection of ookinetes in the mosquito midgut at 26 hpa (Fig. 1A).

**Fig. 1.**
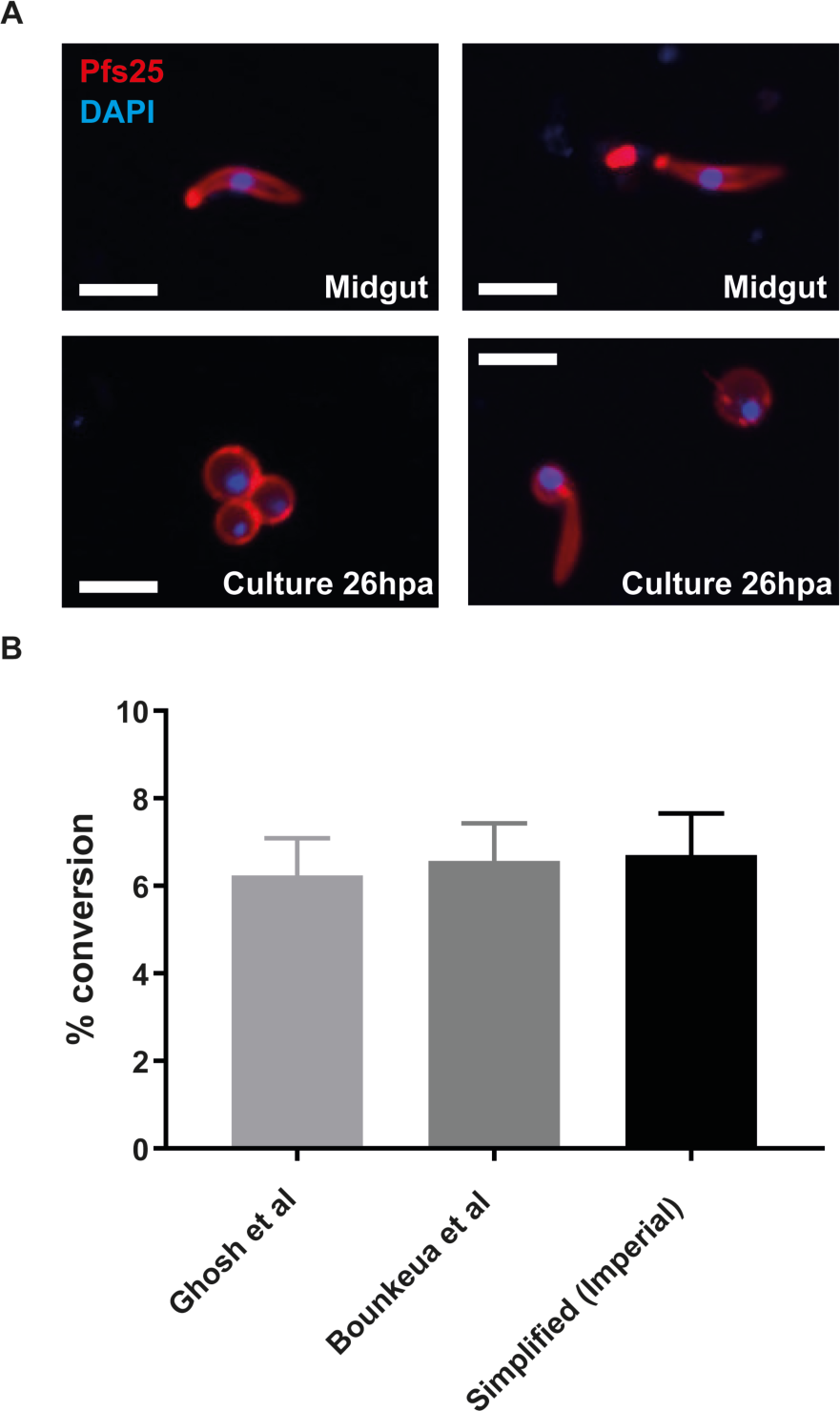
Generation of *Plasmodium falciparum* ookinetes *in vitro* by different methods. (A) Pfs25 is expressed on the surface of female gametes, retorts and ookinetes. Cultured gametocytes were fed to mosquitoes and ookinetes recovered from the midgut at 26 hours post-activation. *In vitro* cultures were induced in parallel and visualized at the same timepoint. Representative images are shown from a minimum of triplicate experiments. Scale bar = 5µm.(B) Two reported *in vitro* ookinete culture protocols (14, 15) and one simplified protocol (defined here) gave similar levels of conversion to retorts (Stage II-III ookinetes); however no fully developed ookinetes were observed. Error bars denote the standard deviation (n = 3 independent experiments).

Using this tool to easily identify and morphologically assess ookinete formation, we compared previously published *in vitro P. falciparum* ookinete culture protocols (see Methods for stage classification). Unexpectedly, in our hands we observed that the majority of Pfs25-surface-positive parasites found *in vitro* were round-shaped (presumed to be unfertilised female gametes). Regardless of the protocol used, the mean percentage of parasites progressing further into retorts (Stages II or III of ookinete development) was not significant between methods – ranging from 6.2% to 6.7% (Fig. 1B, Bonkeua vs. Ghosh p = 0.066). Stage IV retorts - where the nucleus has migrated into the apical protrusion – were seldom observed *in vitro*. Mature ookinetes of stages V and VI were never detected *in vitro,* whilst they were commonly found *in vivo* (Fig. 1A, 2A+B).

### 3.2. Cells reported as cultured ookinetes do not secrete Pfs25 and are recognised by a male-gametocyte specific antibody

Clear morphological differences are apparent between the cells reported as <u>c</u>ultured <u>o</u>okinetes(Bounkeua et al., 2010) (which we term here RCOs) in three previously published reports (Bounkeua et al., 2010; Carter et al., 1987; Ghosh et al., 2010). In our study, Giemsa staining of ookinetes *ex vivo* from mosquito midguts (Fig. 2A) revealed their slim fluke-like morphology with a discrete compact circular pigment-free nucleus, most similar to the RCOs shape described by Carter and colleagues (Carter et al., 1987). In contrast, potential ookinetes found in our cultures (Fig. 2A) had a diffuse nucleus frequently surrounded by pigment, similar to RCOs described more recently (Bounkeua et al., 2010; Ghosh et al., 2010). A further concern with these diffuse nucleus cell types is that they are observed in culture as early as 6 hpa (Fig. 2A) – a timepoint at which midgut ookinetes have yet to form.

**Fig. 2.**
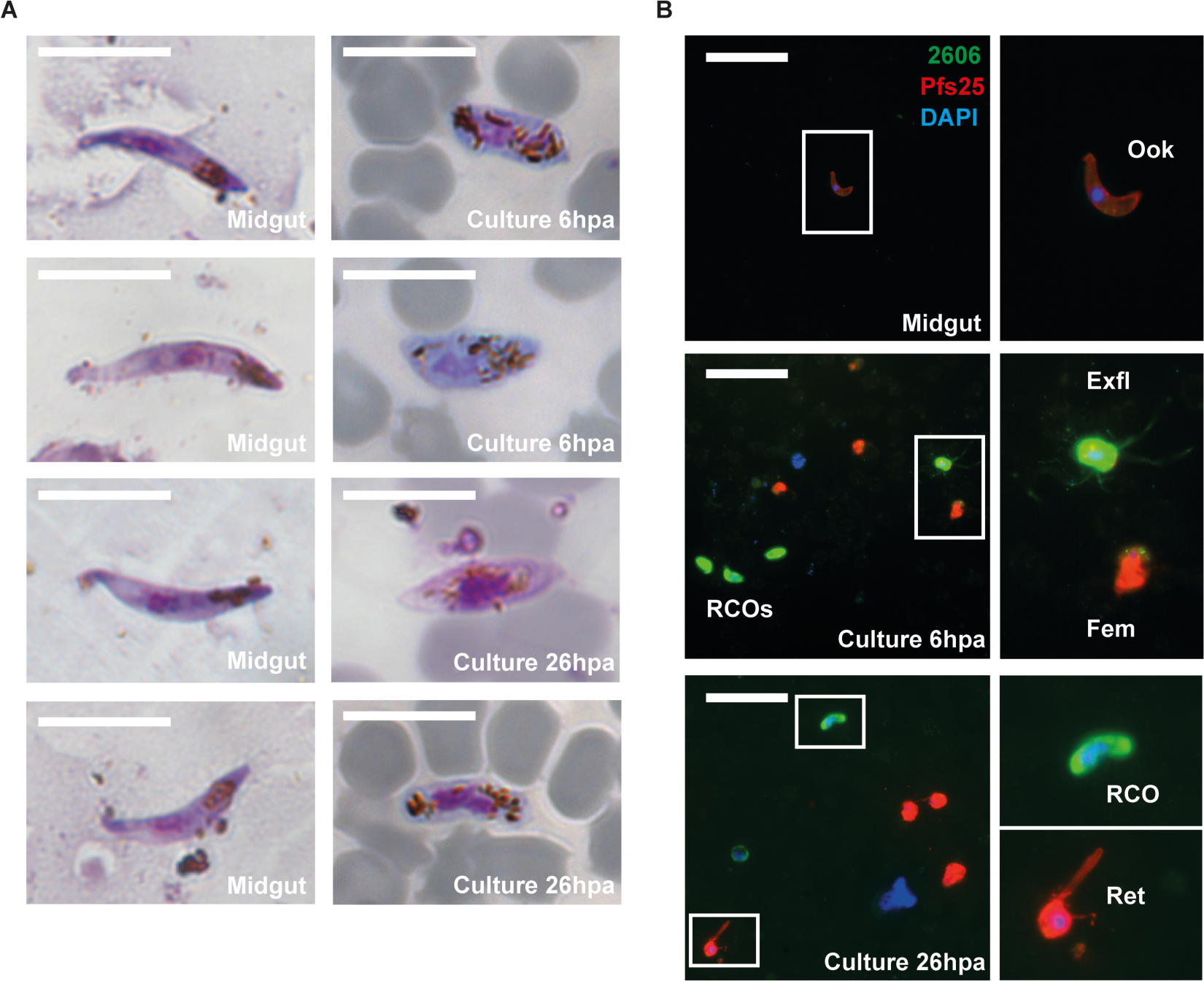
Comparison between *ex vivo* ookinetes and *in vitro* cells reported as ookinetes by morphology and gametocyte sex-specific markers. (A) By Giemsa staining, *P. falciparum* ookinetes isolated from the mosquito midgut show a slim, fluke-like shape, a compact nucleus and a darker apical end (left panels). In contrast, cells previously reported as cultured ookinetes (RCOs) had large elongated nuclei in the centre of the cell surrounded by pigment granules (right panel). Scale bar = 5µm. Representative images are shown from a minimum of triplicate experiment repeats. (B) By immunofluorescence, midgut ookinetes (Ook) express the female gamete/retort/ookinete marker Pfs25, whilst showing very faint staining for the male gametocyte/gamete marker 2606. *In vitro*, female gametes (Fem) and retorts (Ret) were also Pfs25 positive/2606 negative. In contrast, RCOs and exflagellation centres (Exfl) stained strongly for 2606, but not Pfs25. Scale bar = 12µm. Representative images are shown from a minimum of triplicate experimental repeats.

Immunofluorescence showed that these RCO cell types (Fig. 2B) do not express Pfs25 on their surface membrane – the hallmark of female gamete/ookinete identity. In contrast, RCOs, along with exflagellating male gametes and activated male gametocytes, showed strong surface labelling with anti-2606, recognising the male gamete-associated marker PF3D7_1325200. This antibody only very weakly labels midgut ookinetes, and does not stain activated females or retorts (Fig. 2B). Taken together, these morphological and specific antibody data suggest some RCOs are likely to be partially-activated male gametocytes that have replicated their DNA, but have not undergone exflagellation.

In a previous study, electron microscopy sections of some RCOs revealed the presence of apical complexes and microneme containing cells presumed to be ookinetes as these organelles are not present in gametocytes or gametes, although they were not infectious to mosquitoes (Bounkeua et al., 2010). We speculate that these cells are likely arrested at retort stage when the apical complex is present, but not fully mature enough to be infectious.

### 2.3. P. falciparum gametocytes show low fertility in vitro

Given the apparent low rate of gametocyte-to-ookinete/retort conversion that we observed *in vitro*, it was essential to assess the intrinsic *in vivo* viability of our gametocyte cultures. Individual *P. falciparum* gametocyte cultures were divided (n = 6 independent cultures), with half the culture fed to mosquitoes in a Standard Membrane Feeding Assay (SMFA) and the remainder allowed to form ookinetes *in vitro* using our simplified protocol. In parallel, *P. berghei* gametocytes (n = 6 mice) were subjected to a similar SMFA and *in vitro* ookinete culture using the established *P. berghei* protocol. At 26 hpa, cultures were harvested and midgut bloodmeals extracted. All were stained with anti-Pfs25-Cy3 (*P. falciparum*) or anti-Pb28-Cy3 (*P. berghei*) and conversion assessed by quantifying round cells (female gametes) vs retorts/ookinetes. As expected, conversion rates in *P. berghei* were high both *in vivo* and *in vitro* (*in vivo* range: 47.0-57.2%; *in vitro* range: 63.8-83.7%; Fig. 3, Supplementary Fig. 2). *In vitro* conversion was significantly higher, as often seen with *P. berghei*, and is likely attributed to culture medium dilution of inhibitory conditions linked to the high asexual parasitaemia also present. Contrastingly, whilst *P. falciparum* conversion *in vivo* was relatively efficient albeit variable, the corresponding *in vitro* conversion was extremely low (*in vitro* range: 3.2-7.2%; *in vivo* range: 34.9-71.0%; Fig. 3, Supplementary Fig. 2). Taken together, these findings show that our *P. falciparum* gametocytes are viable for onward development in the mosquito; however, unlike *P. berghei*, this does not translate into efficient *in vitro* development. To understand whether this species discrepancy was due to a failure of *P. falciparum* zygotes to progress developmentally, or a failure of male and female gametes to fertilise, we examined parasite fertilisation by flow cytometry. Cultures of *P. falciparum* and *P. berghei* were stained 4-5 hours post activation with anti-Pfs25-Cy3 and anti-Pbs28-Cy3, respectively, as well as Hoechst staining to measure the DNA content within these populations (Fig. 4). Flow analysis detected two discrete populations of wild-type *P. berghei* Pbs28-surface-positive cells, with different Hoechst intensities (Fig. 4A). Using a *P. berghei* mutant line (PbGEST-KO) that only rarely fertilises for comparison (Talman et al., 2011), the lower-intensity Hoechst peak was inferred to be unfertilised Pbs28-positive female gametes. The higher-intensity peak observed in the WT parasites was virtually absent from the PbGEST-KO, suggesting this peak represents successful fertilisation events. Supporting this, the percentage of cells in this peak closely matches the *P. berghei* conversion rate observed *in vitro* (Fig. 3). In *P. falciparum*, two populations of Pfs25-positive cells were observed – one forming a large peak at low Hoechst intensity and the other forming a very small broad peak at higher Hoechst intensity (Fig. 4B). Again, the higher-intensity peak approximates to the conversion rate we observed by microscopy for *P. falciparum* (Fig. 3) and resembles the pattern of the PbGEST-KO. Taken together, these data supports the hypothesis that the low efficiency of *in vitro P. falciparum* ookinete culture can largely be accounted for by failure of the gametes to fertilise; however, clearly those cells that do fertilise do not mature into fully formed ookinetes and so other factors may also contribute.

**Fig. 3.**
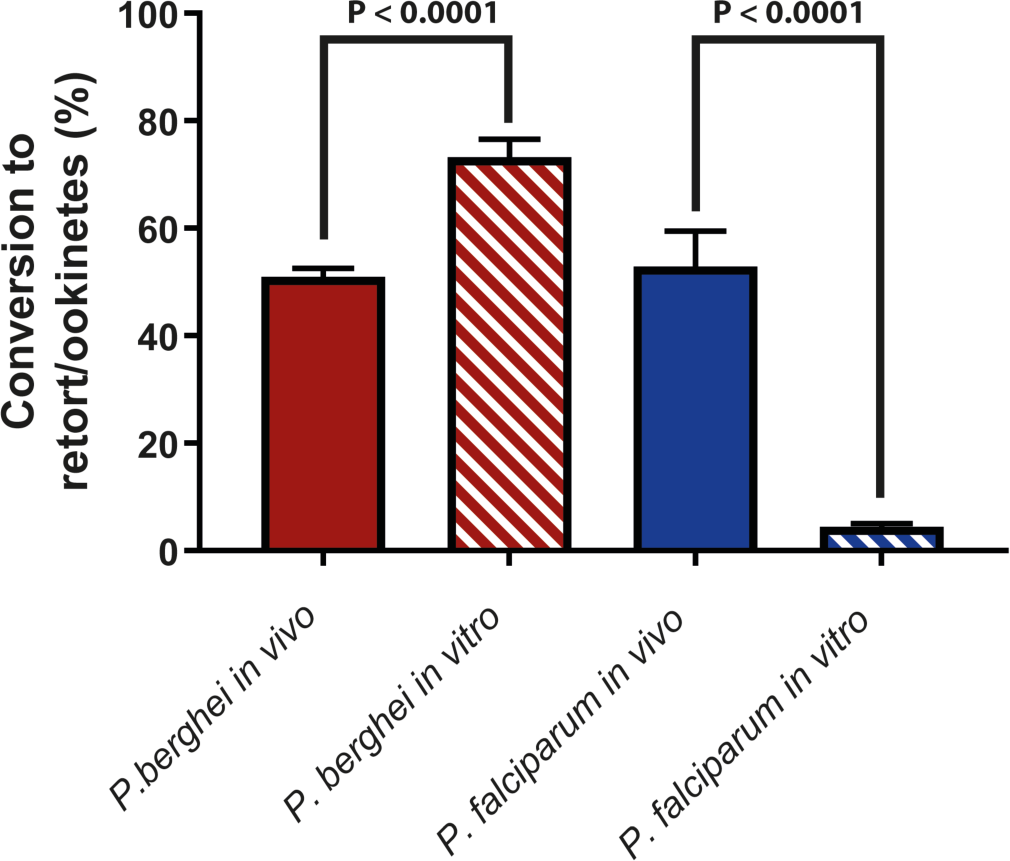
Fertility of *Plasmodium berghei* and *P. falciparum* gametocytes *in vitro* and *in vivo*. *P. berghei* (red bars) and *P. falciparum* (blue bars) gametocytes were induced to develop ookinetes *in vitro* (hatched bars). In parallel, the same parasite samples were also fed to mosquitoes to assess their intrinsic fertility *in vivo* (solid bars). At 26 hours post-activation, parasites were stained live with Cy3-conjugated antibodies specific for Pfs25 (*P. falciparum*) and Pbs28 (*P. berghei*). Conversion rates were determined as the percentage of retort/ookinete-shaped cells to round cells (unfertilised females). n = 6 paired experiments. Error bars denote the standard deviation. Significance determined by T-test.

**Fig. 4.**
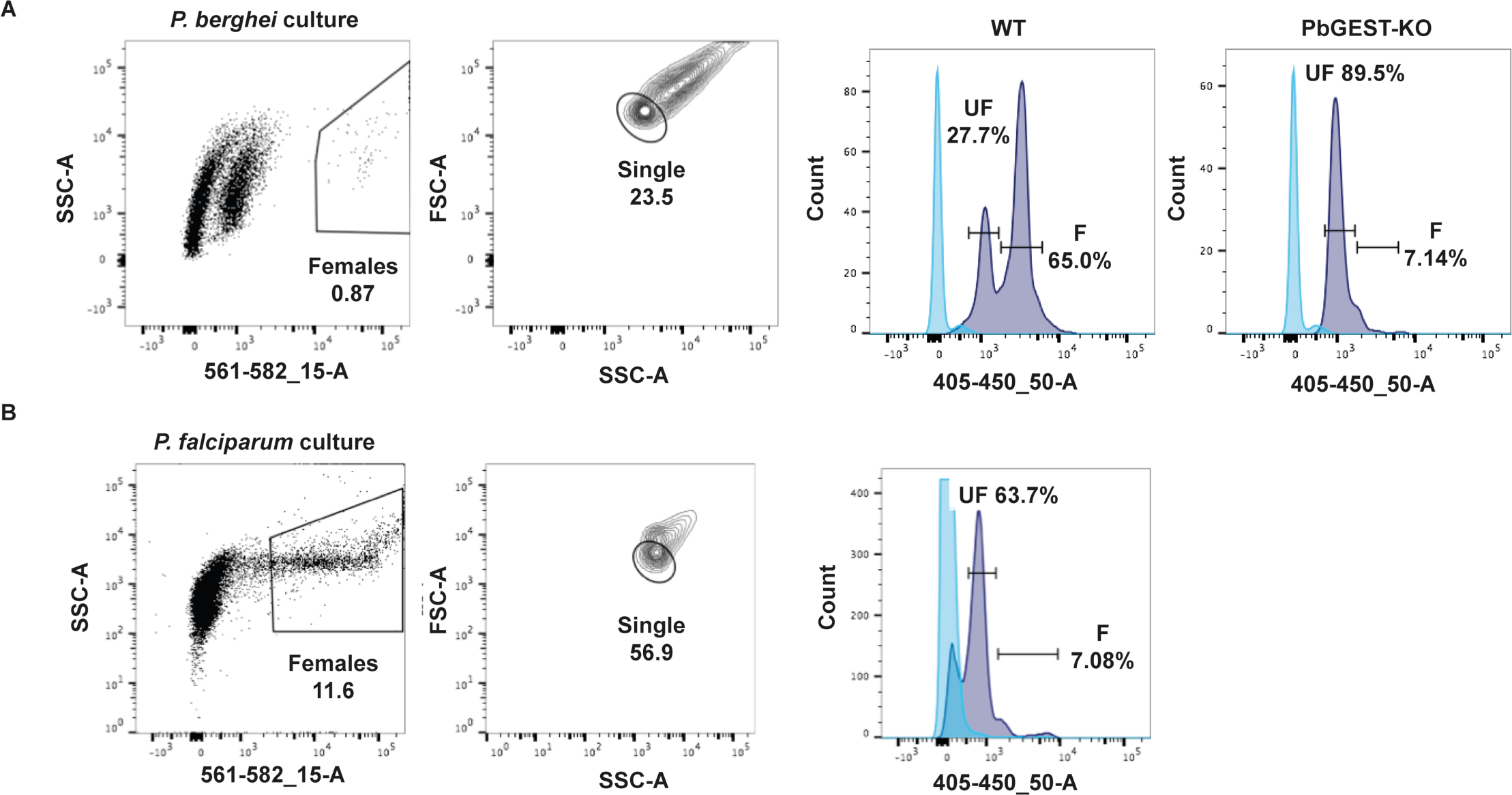
DNA content suggests *Plasmodium falciparum* gametocytes poorly fertilise *in vitro*. Representative flow cell sorting profiles of parasite cultures from a minimum of triplicate experimental repeats. Females were detected with 13.1-Cy3 anti-Pbs28 in *P. berghei* (A) and 4B7-Cy3 anti-Pfs25 in *P. falciparum* (B). Dot plots display cell subpopulations based on side-scattered light (SSC) and Cy3 intensity (561-562_15-A); the Cy3 surface positive female population is gated. Contour plots display subpopulations based on SSC and forward-scattered light (FSC), singlets (si) were selected within the female population specifically to exclude aggregated gametes, zygotes, retorts and ookinetes which adhere robustly to each other (31). Histograms represent the number of female parasites with a given Hoechst (405-450_50-A) profile. Light-blue represents Hoechst-unstained female parasites (background). Dark blue represents Hoechst stained female parasites. Fertilization events in wild-type *P. berghei* (WT) occur at high frequency but are rare in GEST knockouts (PbGEST-KO) (17), which display a high number of unfertilized events. Fertilization events in *P. falciparum* are low and most similar to the profile of *P. berghei* GESTKO.

### 2.4. Systematic modifications to culture protocols towards improved fertilisation/ookinete yield

Multiple factors could contribute the inefficient development of *P. falciparum* ookinetes *in vitro*. Using a systematic literature-based approach, we tested a range of modifications to our simplified culture protocol and evaluated their effect on conversion rate. Modifications broadly fell into three categories (Fig. 5, Supplementary Tables 1+2): 1. Parasite factors; 2. Additives to gametocyte culture; 3. Ookinete culture conditions. In all experiments, modifications were evaluated compared to our unmodified protocol. Substantial experimental variation in conversion rate between control experiments was observed (Fig. 5A). The two most widely used *P. falciparum* strains for transmission studies are NF54 (used here) and 3D7 – a clone originally derived from NF54. In a head-to-head comparison, NF54 gave a significantly higher mean conversion rate than 3D7 (8.0% vs 3.1%, respectively; Fig. 5B), justifying its use in this study. To understand if gametocyte maturity could be linked to *in vitro* fertility, gametocyte cultures of ages routinely reported in the literature as used for mosquito infections (Lensen et al., 1999; Miura et al., 2013) were tested for ookinete conversion (Fig. 5C). Cultures of 14 to 18 days old (and various mixed combinations) showed no significant differences in conversion efficiency.

**Fig. 5.**
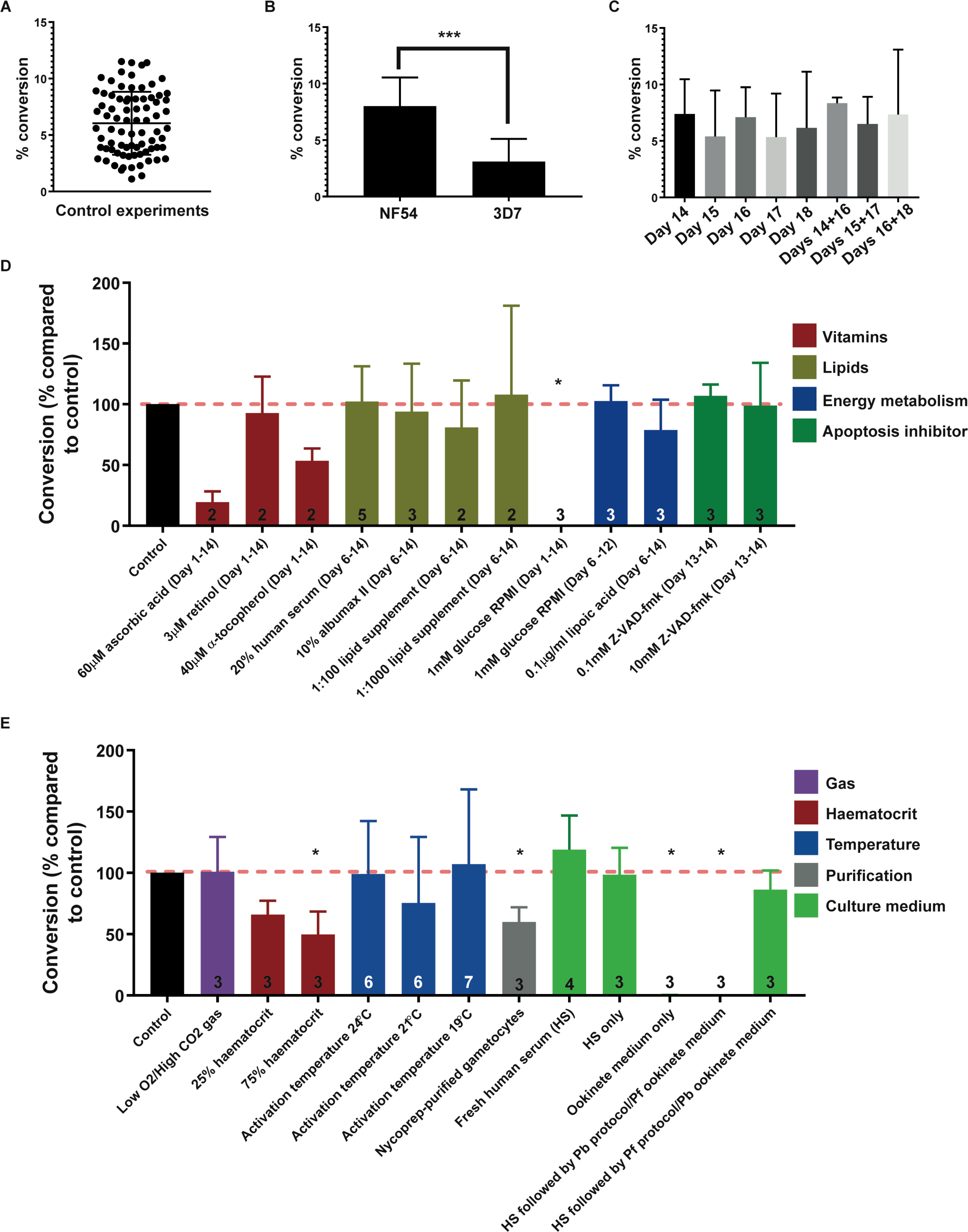
Investigating factors affecting the efficiency of *Plasmodium falciparum* ookinete conversion. To better understand factors critical for P. falciparum *in vitro* ookinete culture, we tested a wide range of conditions and assessed conversion rate. (A) Each experiment was compared with reference to our basic control protocol as conversion rates varied substantially between different gametocyte preparations. Each datapoint is an independent control culture (n = 79), line indicates the mean conversion rate and error bars denote the standard deviation.(B) *P. falciparum* NF54 strain was found to be significantly more efficient at fertilisation than 3D7 strain (P < 0.001, unpaired student’s T-Test). (C) The age of the culture had no impact on gametocyte fertility. (D) The impact of various factors on gametocyte fertility added to the developing gametocyte culture (pre-activation factors). Numbers on bars indicate the number of independent replicates performed and error bars denote the standard deviation. * indicates P < 0.05, unpaired student’s T-Test. (E) The impact of various factors on gametocyte fertility added to the parasites after ookinete culture induction (post-activation factors). Numbers on bars indicate the number of independent replicates performed and error bars denote the standard deviation. * indicates P < 0.05, unpaired student’s t-test.

One of the key differences between *P. falciparum* and *P. berghei* gametocytes used in these conversion experiments is that the *P. berghei* gametocytes have developed and matured within their mammalian host rather than artificial culture conditions. Therefore, it was speculated that the *in vitro* fertility of cultured gametocytes might be improved by addition of “missing” *in vivo* factors during culture development (Fig. 5D). Vitamins absent from RPMI medium formulations were added throughout the gametocyte culture period, but had no statistically significant effects on conversion rate. Similarly supplementing or replacing the standard 10% human serum found in our standard culture medium with other lipid sources specifically during gametocyte development (rather than asexual growth before gametocyte commitment) gave no noticeable improvement. Previous metabolomic analyses suggest that gametocytes differ in mechanisms of energy metabolism compared to asexual parasites (Lamour et al., 2014; MacRae et al., 2013). It was found that gametocyte development (Days 6-12) and subsequent ookinete conversion were not affected (certainly not improved) by low glucose (1mM) RPMI. In contrast, low glucose could not support gametocyte culture growth at all if also present during the asexual culture phase (Days 1-14). Gametocytes were pre-treated for 24hr before induction (and during ookinete culture) with the apoptosis inhibitor Z-VAD-fmk, which has previously been shown to enhance parasite survival in the mosquito in *P. berghei* (Al-Olayan et al., 2002). No appreciable difference was observed, indicating parasite apoptosis is not a major factor in fertility.

Since the conditions tested here for pre-ookinete culture induction did not have a positive impact on conversion efficiency, alterations to conditions post-induction were tested (Fig. 5E). *berghei* ookinete culture is routinely performed at ambient gas concentrations. We found that culturing in a low oxygen, high carbon dioxide environment (asexual “malaria gas”) had no impact on fertility. Varying the haematocrit of the initial induction mix in human serum from 50% to 25% or up to 75% reduced conversion efficiency. A range of incubation temperatures also showed no impact. To promote closer interaction between male and female gametocytes (hence increase the chances of gamete encounters) gametocytes were purified away from the erythrocytes comprising the bulk of the gametocyte culture using Nycoprep density centrifugation (Lelièvre et al., 2012). Under these conditions, conversion rates were significantly reduced (P < 0.05), suggesting either cell proximity is not a critical factor or that purification is detrimental to gametocyte fertility. Our simplified protocol involves reuspending the pelleted gametocyte culture in human serum to 50% haematocrit for 30 minutes to activate, form gametes and fertilise. Afterward, Pf ookinete medium is added to maintain the developing cells. To better understand which elements of the culture protocol were most important, different variations were tested. The initial human serum incubation at 50% haematocrit (activation/fertilisation step) was found to be essential and sufficient for conversion rates identical to our control protocol, and could not be replaced with ookinete medium alone, which totally abrogated conversion. *P. falciparum* ookinete medium formulation could be replaced with *P. berghei* ookinete medium (containing xanthurenic acid) provided the initial human serum incubation step was maintained.

This extensive assessment of variables strongly suggests that the single most critical factor for *P. falciparum in vitro* gamete fertilisation, even at a far lower rate than *in vivo*, appears to be the initial gamete formation step in human serum. However, those parasites that are fertilised are still not able to continue development *in vitro* to fully mature ookinetes.

## 4. Discussion

Successful and efficient *in vitro* generation of *P. falciparum* ookinetes is a major roadblock to malaria transmission research in the laboratory. Our data shows that there is nothing intrinsically wrong with the fertility of cultured *P. falciparum* gametocytes as they successfully infect mosquitoes. However, *in vitro* gamete fertilisation during the initial period of ookinete culture is highly inefficient, and those that do fertilise do not fully mature into Stage V ookinetes. It also appears that high concentrations of human serum is essential to observe any fertilisation. When considered together, these data suggest that *P. falciparum* ookinete culture is either lacking a factor critical to generating fertile gametes, or *in vitro* conditions introduce an inhibitory factor that prevents successful fertilisation.

Our data also suggests caution when identifying *in vitro* cultured ookinetes. To this end, we suggest the following criteria that must be met to definitively prove the ookinete identity of cultured cells: 1. Appropriate morphological comparison with mosquito-derived ookinetes in parallel experiments; 2. Surface expression of Pfs25 (and other known ookinete proteins such as CTRP if possible); 3. Observed *in vitro* motility through a matrigel matrix (Moon et al., 2009); 4. Infectiousness of cultured ookinetes to mosquitoes.

Whilst ultimately unsuccessful in our goal to generate *in vitro P. falciparum* ookinetes, we hope our investigation highlights areas for further investigation and direct future studies.

## Supplementary Material

### Supplementary File 1

This file contains the following: Supplementary Fig. 1. Characterization of anti-2606 antibody against PF3D7_132520; Supplementary Fig. 2. Paired *in vivo*/*in vitro* ookinete conversion – individual replicates; Supplementary Table 1. Parameters tested to improve gametocyte culture (pre-activation); Supplementary Table 2. Parameters tested to improve ookinete culture (post-activation).

## Acknowledgements

We acknowledge the large number of researchers worldwide that have trialled various methods towards *P. falciparum* ookinete culture and shared thoughts at conferences, meetings and directly through email in helping this study. We thank Mark Tunnicliff for technical assistance with mosquito rearing. We acknowledge MR4, part of the BEI Resources Repository, for provision of *Plasmodium falciparum* anti-Pfs25 MAb 4B7 (MRA-28) deposited by D. C. Kaslow. We would like to thank VigiLab (Imperial College London) for providing us with *Anopheles coluzzii* and particularly Kathrin Witmer for fruitful discussions. We also thank Catherine Simpson from Imperial College Flow Cytometry Facility for training and support. This work was supported by the Wellcome Trust [100993] Investigator Award to JB, [088381] Research Career Development Fellowship to NG, and Institutional Strategic Support Fund Inflammation Science Studentship to JG; Bill & Melinda Gates Foundation, grant OPP1043501 (MJD, RES and JB); Biotechnology and Biological Sciences Research Council doctoral training program studentship to SS.

